# Reading Acquisition Drives Linguistic Cross-Modal Convergence in The Left Ventral Occipitotemporal Cortex

**DOI:** 10.1101/2024.11.26.625405

**Authors:** A Dębska, S Wang, K Jednoróg, C Pattamadilok

**Affiliations:** Laboratory of Language Neurobiology, Nencki Institute of Experimental Biology, Polish Academy of Sciences.; Aix Marseille Univ, CNRS, LPL, Aix-en-Provence, France

**Keywords:** reading acquisition, left ventral occipitotemporal cortex, cross-modal integration, print-speech convergence

## Abstract

Reading acquisition requires linking visual symbols with speech sounds, leading to the development of neural sensitivity to print. While prior studies have shown the importance of cross-modal integration in spoken language areas, the higher-level visual area (lvOT) processing printed words remained more context-dependent. This longitudinal study investigated whether the lvOT undergoes cross-modal reorganization to facilitate print-speech integration during reading development and how these changes relate to reading skills. We followed children over two years, beginning at the onset of formal reading instruction. We examine lvOT responses to print-specific, speech-specific, and its convergence at whole-brain, region of interest and voxel-based levels. Results showed that with reading experience, the Initial print-specific responses in the lvOT are transformed into responses to both print and speech input. This transformation positively correlates with reading skills, especially in early acquisition stages. These findings suggest that reading acquisition drives cross-modal reorganization within the lvOT, enabling the area to integrate print and speech. They shed light on the broader neural mechanisms supporting reading development.

## INTRODUCTION

The fundamental stage of reading acquisition is learning to associate abstract visual symbols with speech sounds. This learning process gives rise to a novel functional role in subpopulations of neurons in the left ventral occipitotemporal cortex (lvOT), enabling the recognition of written words (Brem et al., 2010). Following the assumption that the new functional response in these neurons is “print-specific”, i.e., showing preferential responses to known written words over other visual categories (Dehaene & Cohen, 2011), the area has been called the “Visual Word Form Area” (VWFA, Cohen et al. 2000, 2002). In previous studies, print-specificity (stronger responses to words compared to false fonts) in the lvOT emerged in pre-reading children already after less than 4 hours of laboratory-based grapho-phonological training (Brem et al., 2010). In more ecological classroom settings, this activation was observed in beginning readers shortly after the onset of formal reading instruction (Chyl et al., 2018, Dehaene-Lambertz, 2018). Crucially, print-specific activity has been reported only when visual symbols are associated with speech sounds, not with non-speech inputs like tones or noise (Hashimoto & Sakai, 2004). This observation suggests that this response pattern is restricted to the processing of linguistic materials.

Another important, though less investigated, functional reorganization in the lvOT is the increased of sensitivity to cross-modal language input (Dehaene et al., 2010, Rueckl et al., 2015, Preston et al., 2016, Marks et al. 2019, McNorgan & Booth, 2015, Chyl et al., 2018, 2021, Dębska et al., 2019, Planton et al., 2019). The universality of the reading network across language systems is explained by the proposal that the reading network emerged from the phylogenetically older, pre-existing spoken language and non-language visual networks (Frost et al., 2012, Rueckl et al., 2015, Chyl et al., 2021). To date, the most common evidence for cross-modal reorganization is *print-speech convergence*, which reflects joint activation to both print and speech stimuli within the same brain region. These convergent responses and their relationship with reading ability are established phenomena in frontal and auditory cortices within the spoken language network (Frost, 2009; Marks et al., 2019; Preston, 2016; Rueckl, 2015; Chyl et al., 2018, 2021, McNorgan et al., 2014). The most common convergence zones include the bilateral superior temporal gyri (STG), middle temporal gyri (MTG), and the left inferior frontal gyrus (IFG). The print-speech convergence in those areas has been linked to the reading level already in beginning readers (Preston et al., 2016, Chyl et al., 2018). Additionally, the level of print-speech convergence in the bilateral STG, MTG, and IFG differentiated typical readers form dyslexic readers (Yan et al., 2024, Dębska et al., 2021).

While the convergence of print and speech in spoken language-related brain areas like the STG, MTG, and IFG is well-documented and linked to reading ability, evidence for a similar reorganization in the lvOT remains limited, inconsistent across languages and points to the context-dependent rather than a universal role of cross-modal reorganization in this brain region. Indeed, previous studies have shown print-speech convergence in the lvOT in an opaque English and Hebrew orthography (Preston et al., 2016, Rueckl et al., 2015) but not in more transparent Polish and Spanish (Dębska et al., 2019, Chyl et al., 2021, Rueckl et al., 2015), suggesting that this reorganization may depend on orthographic transparency which reflects the strength of the connections between spellings and sounds (Katz & Frost, 1992).

Additionally, previous findings often lack comparisons to non-linguistic conditions, making it challenging to isolate language-specific convergence from visual-auditory multimodal convergence (see e.g., Dzięgiel et al., 2021, McNorgan & Booth, 2015). A previous study that focused on linguistic print-speech specific convergence failed to find effects within lvOT even in an opaque English orthography, while more robust evidence was found in the MTG or STG regardless of the transparency of orthography and the choice of control conditions (Chyl et al., 2021).

Most findings on print-speech convergence in the lvOT predominantly came from cross-sectional studies, making it challenging to understand the developmental trajectory of cross-modal reorganization in the lvOT and to capture the progression of functional changes over time. To our knowledge, only one longitudinal study was conducted on young children (Preston et al., 2016), tracking participants from 8.5 to 10 years old, who had already mastered reading to some extent. The present study followed the same line of longitudinal protocol while focusing on the earliest and critical period of reading acquisition where children started to learn letter-sound associations and two years later when reading had become more automatized.

### Current study

The present study aimed to determine whether reading acquisition induced a functional reorganization within the lvOT that allowed an integration of responses to print and speech, as reported for other spoken language regions. To this aim, we used longitudinal protocol in which a group of young children were followed from the beginning of formal reading instruction (Mean age=6.9 years old) until two years later. This protocol allowed us to track the earliest stage of the development of the lvOT’s specific responses to print (compared to symbols, hereafter referred to as *print-specific processing*), to speech sounds (compared to vocoded speech, hereafter referred to as *speech-specific processing*), and, most importantly, the development of joint responses to print and speech-specific processing, which reflects the *print-speech specific convergence*.

We hypothesized that the lvOT’s sensitivity to speech-specific sounds and the convergence between print and speech-specific stimuli would increase with reading ability. To complement existing studies, we examined this phenomenon in detail at three complementary spatial scales. First, at the whole brain level, to obtain a global overview of print-specific network, speech-specific network and their conjunction reflecting print-speech specific convergence, at the beginning of formal reading instruction and two years later. Second, we analyzed response patterns to print and speech-specific stimuli and their conjunction with the relation to reading ability at the lvOT cluster level. Finally, within this ROI, we investigated the development of print-speech specific convergence by analyzing changes in functional responses to written and spoken input within individual lvOT voxels across two time points, and explored how these changes were related to children’s reading scores.

## METHODS

### Participants

All participants took part in a longitudinal study on reading development and dyslexia approved by the Warsaw University Ethical Committee and conducted in accordance with the provisions of the World Medical Association Declaration of Helsinki. Parents of participating children signed informed consent forms and the children gave verbal assent.

Behavioral data were collected at three time points with one-year intervals. However, the fMRI data were collected only at the first and last time points. For clarity, we refer to these two fMRI time points as T1 and T2.

We recruited a total of 120 native Polish-speaking children from first-grade and kindergarten that met the inclusion criteria: 1) Typical or higher IQs (above the 25th percentile as measured with Raven’s Matrices); 2) Polish monolingual; 3) right-handed, as reported by parents; 4) born at term (≥37 weeks of gestation); 5) no history of neurological illness or brain damage and 6) no prior symptoms of ADHD as reported by parents.

The successful scanning at T1 included 92 children (see Dębska et al., 2016). T1 and T2 sessions were conducted on 90 children (see Łuniewska et al., 2019). In the current study, we focused on children who demonstrated typical reading development. Therefore, the analyses were conducted on the data from 68 children (see Table 1 for demographics) who were not diagnosed with reading deficits at T2, based on the standardized battery for dyslexia diagnosis in Poland (Bogdanowicz et al. 2008). At T1, 20 participants were in their last year of kindergarten (Mean age = 6.88, SD = 0.66) and 48 were first graders (Mean age = 7.01, SD = 0.48). Consequently, at T2, there were 20 second-graders and 48 third graders.

**Table 1.** Averaged ages and performances obtained in the behavioral tasks by the group of 68 children. Standard deviations are in the parentheses.

|  | Age T1 | Age T2 | Nonverbal IQ* | Letter T1** | Word T1*** | Orthography task T1# | Word T2 |
| --- | --- | --- | --- | --- | --- | --- | --- |
| Mean | 6.9 (0.5) | 8.9 (0.5) | 78.6 (17.1) | 51.4 (15.3) | 20 (19.2) | 20(3.7) | 76 (22) |
| Min | 5.8 | 7.8 | 24 | 4 | 0 | 16 | 43 |
| Max | 8.1 | 10.1 | 98 | 64 | 69 | 29 | 125 |

\*Standardized score, *Raven Matrices Test*, measured only at T1; \*\* Knowledge of letters, maximum score = 64 (upper and lower cases); \*\*\* Number of words read correctly in 60 seconds; # maximum score = 30.

Given the educational reform unfolding in Poland during our study, parents could choose to start their children’s formal schooling at the age of six or seven. It is noteworthy that, although the formal onset of reading instruction typically begins in elementary school, the kindergarten curriculum already includes preliminary letter instruction. In our pool of participants, some children were also taught to read by their parents. Our reading assessments showed that, at T1, 12 out of 68 children could not read any word from our reading test (0 words per minute). However, they were able to name letters and perform the orthographic task (described below) to some extent, which suggested that even with limited exposure to written material at that age, they had letter knowledge and demonstrated some sensitivity to the graphotactic properties of the script (see Table 1SM).

### Behavioral tasks

Following the fMRI session (described below), children were tested with a battery of behavioral tasks. The results are described in Table 1. In the letter knowledge task, children were asked to name all letters of the Polish alphabet presented in upper and lower case (max score = 64: 32 upper and 32 lower case letters). In the word reading task, children were asked to read aloud as many words as possible in 60 seconds (score: number of words read correctly per minute, maximum score = 150; Szczerbiński & Pelc-Pękała, 2013). In the orthography task children were presented with 30 pairs of letter strings (trigraphs: e.g. DAG - DGA, Awramiuk et al., 2013) and, for each pair, they had to select the item that resembled Polish orthography. Nonverbal IQ was measured with a standardized version of the Raven’s Colored Progressive Matrices (Szustrowa et al., 2003).

### Fmri task

fMRI data were collected using a 3T Siemens Trio scanner equipped with a 12-channel head coil, employing a whole-brain echo-planar imaging sequence (32 slices, 4 mm slice thickness, TR = 2000 ms, TE = 30 ms, flip angle = 80°, FOV = 220 mm², matrix size: 64 × 64, voxel size: 3 × 3 × 4 mm). Anatomical images were captured with a T1-weighted sequence (176 slices, 1 mm slice thickness, TR = 2530 ms, TE = 3.32 ms, flip angle = 7°, matrix size: 256 × 256, voxel size: 1 × 1 × 1 mm). Prior to scanning, children were familiarized with the experimental setup and the MRI environment using a mock scanner. During the fMRI session, participants were exposed to print and speech stimuli representing four conditions: (1) printed words, (2) spoken words, (3) symbol strings, and (4) vocoded speech. The printed and spoken words were high-frequency, short Polish words from the Polish CHILDES database (Haman et al., 2015). For the print condition, the average word length was 4.16 letters (SD = 0.86), and for the speech condition, it was 4.14 letters (SD = 0.85). The average number of phonemes was 3.85 (SD = 0.75) for both conditions. Words were matched on parameters such as letter count, phoneme count, syllable count, and frequency (see Supplementary Materials, S1, Chyl et al., 2018). Symbol strings were created by replacing each letter in the printed word list with a corresponding symbol from the Wingdings font. Vocoded speech stimuli were generated from the spoken word list using Praat software (Boersma and Weenink, 2001), which divided the speech signal into three frequency bands, applied the dynamic amplitude contour to noise, and recombined the signals, preserving the temporal structure but distorting the phonetic content. Auditory stimuli were played through MRI-compatible noise-canceling headphones (CRS) at approximately 70 dB, while visual stimuli were displayed on a 32’’ LCD screen (70 x 40 cm) in black text/symbols on a white background.

Each condition featured 96 stimuli, organized into 24 trials consisting of four stimuli each. Visual stimuli were shown for 250 ms, followed by a 200 ms blank screen, and auditory stimuli lasted 800 ms. The experiment used “jittered” intertrial intervals with occasional null trials (a black fixation cross on an empty screen), leading to ITIs ranging from 4 to 13 s (average: 6.25 s). A total of 96 trials, divided between two runs, were presented, with each run featuring 12 trials from each condition. This resulted in 48 trials per run (4 conditions × 12 trials), which were shown in a pseudorandomized sequence with the constraint of no more than two consecutive trials from the same condition. Stimuli were presented using Presentation software (Neurobehavioral Systems, Albany, CA).

Children were instructed to attend to the stimuli, aiming to rapidly activate the language network in each modality by comparing the non-linguistic and linguistic input. The task has been validated in previous fMRI and EEG studies examining print and speech sensitivity in children (Malins et al. 2016; Chyl et al. 2018, 2019a, b; Lochy et al. 2015, Dębska et al., 2019).

### Fmri data processing and first-level analysis

The neuroimaging data preprocessing and analyses were performed using Statistical Parametric Mapping (SPM12, Welcome Trust Centre for Neuroimaging, London, UK) run on MATLAB R2016b (The Math-Works Inc. Natick, Massachusetts, United States).

Images collected at the two time points were realigned to the mean. Next, a pairwise longitudinal registration was performed on T1-weighted images from two time points and a midpoint average image was created. The outcome of the pairwise longitudinal registration was coregistered to the mean functional image. Coregistered images were segmented using pediatric tissue probability maps, while the Template-O-Matic toolbox (Wilke et al., 2008) was used with the matched pairs option. The functional images were normalized using compositions of flow fields and a group-specific template. The normalized functional images were smoothed with an 8 mm isotropic Gaussian kernel. Finally, the functional images were temporally normalized to percent signal change in each voxel (Chen et al., 2017; Schilling et al., 2023). The data were modeled for each run using the canonical hemodynamic response function convolved with the tasks. ART toolbox was used to reject motion-affected volumes surpassing the movement threshold of 3 mm and a rotation threshold of 0.05 radians (Dębska et al., 2019, 2021, 2023; Raschle et al., 2012). Subjects were included if a minimum of 80% of volumes from each run were artifact-free. All subjects in the group fulfill this criterion.

A random effect GLM was computed for each participant and condition. We conducted a subject-level analysis of the experimental conditions: print, speech, symbol strings, vocoded speech including six motion parameters and separate regressors for each volume identified as motion-affected by ART toolbox. In the first-level analysis, contrasts were generated for each condition at T1 and T2 against baseline (rest) and between conditions (*print-specific*: print > symbol strings; *speech-specific*: speech > vocoded-speech). All models were run with an explicit grey matter mask based on the pediatric template created to match the age of our sample (Template-O-Matic toolbox, Wilke et al., 2008).

### Whole-brain analysis

To identify the *print-specific* and *speech-specific* language networks, we conducted one-sample t-tests on the print > symbol strings and speech>vocoded one-sample t-tests for contrasts at T1 and T2 separately. We also calculated a logical conjunction map between print-specific and speech-specific contrasts to localize significant clusters of co-activation between print and speech. Group conjunctions were based on significantly active voxels at *p* < 0.001, FWEcc in both print > symbol strings and speech > vocoded contrasts. All results are presented with the correction for multiple comparisons with the primary threshold p<0.001 (voxelwise) and p<0.05 FWE cluster correction. To compare our results with previous studies we also performed the same analysis based on activity in print-rest and speech-rest contrasts (see SI xxx).

### The left-vOT analysis

The second set of analyses focused on the lvOT (see Figure 1). We restricted our analysis to an area defined by an anatomical mask of the left Fusiform Gyrus and Inferior Temporal gyrus (AAL atlas, Tzourio-Mazoyer et al., 2002). The mask was also restricted by the grey matter mask pediatric template that matched the age of our sample. The mask included 2189 voxels with MNI center coordinates of −52 −54 −28 and y-axis from −32 to −74. This is analogical to Bouhali et al., (2019) definition on the y-axis side (−30 to −70) in the study on functional and structural segregation of lvOT. This allowed us to track the development of brain responses to visual and auditory stimuli within the posterior and anterior parts of the lvOT, which show differential function in print and speech processing in reading development (Dębska et al., 2023, Wang et al., 2018, 2021).

**Figure 1.**
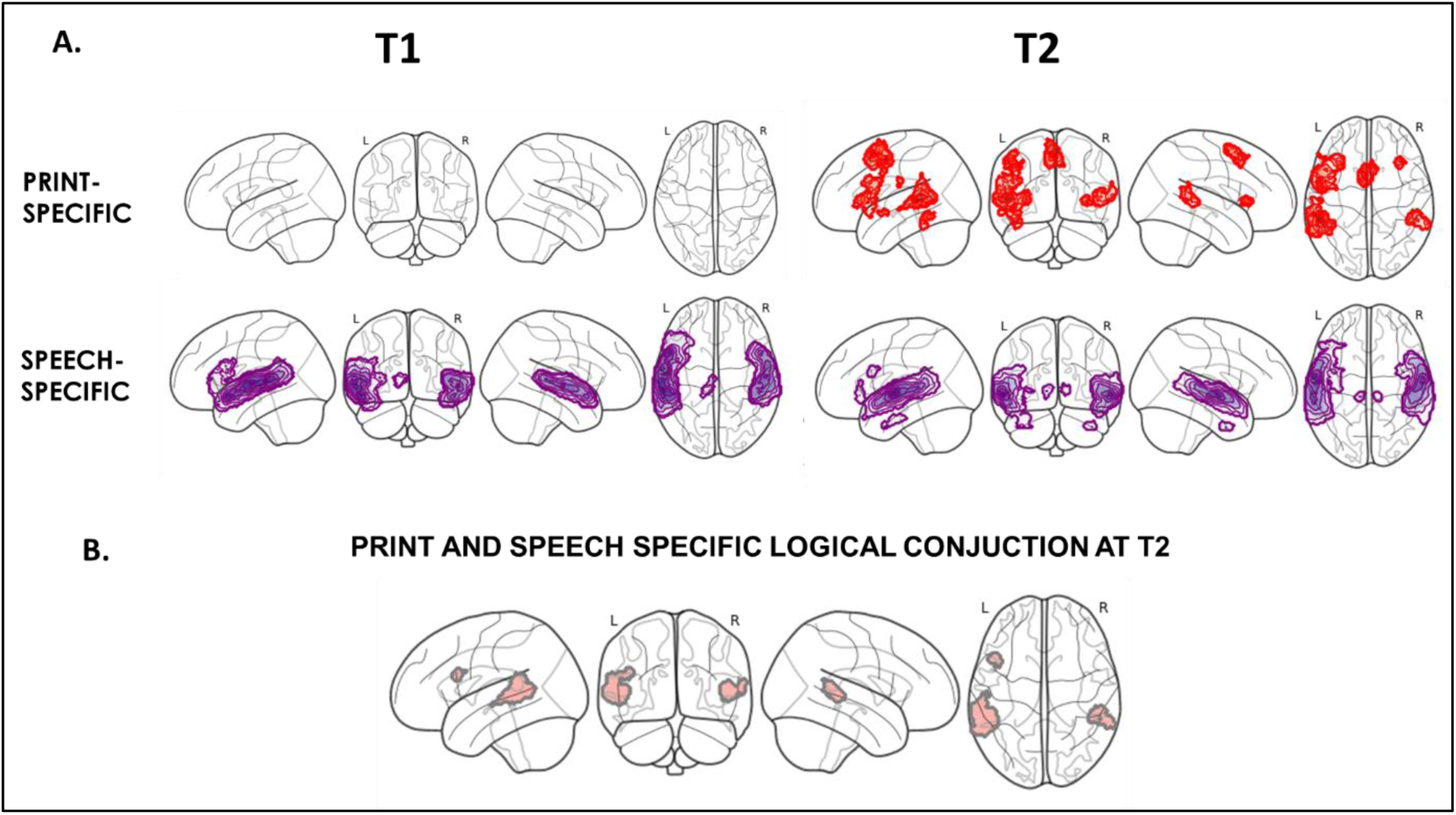
A. One-sample t-tests for print-specific (print>symbol strings, red) and speech-specific (speech> vocoded speech, purple) activation at the first (T1) and second time-points (T2). B. Logical conjunction results at T2

As we did not obtain print-speech specific convergence maps at *p* < 0.001, FWEcc within the lvOT mask, we focused on the activation from print-specific and speech-specific contrasts. We ran 2 (print-specific, speech-specific) x 2 (T1, T2) repeated measure ANOVA within the lvOT mask.

To examine the relationship between brain activity and reading skills, for each type of contrast (print-specific and speech-specific) and each time-point (T1 and T2), we performed a multiple regression model for brain activity within the left-vOT mask and reading score, with Age, Sex, Non-verbal IQ as covariates. To ensure that the subgroup of children with the lowest reading scores did not drive the results, an additional regression model was also conducted on a restricted group of children excluding those with a score of 0 in the word reading task at T1 (N=12, see: SM1).

All results are presented with the correction for multiple comparisons FWE with the primary threshold p<0.001 (voxelwise) and p<0.05 for a cluster.

### The individual voxels analysis within the left-vOT

We examined the developmental changes in print-specific and speech-specific responses in lvOT in greater detail. Within the lvOT mask, based on the 1^st^ level activation values for each participant, we defined four categories of voxels: 1) **Print- and Speech-specific voxels** (i.e., the voxels active above threshold where beta value β > 0 in both print-specific and speech-specific contrasts; 2) **Print-specific voxels** (i.e., the voxels active above threshold beta value β > 0 in print specific contrast and β < 0 in speech specific contrast); 3) **Speech-specific voxels** (i.e., the voxels active above threshold beta value β > 0 in speech specific contrast and β < 0 in print specific contrast); 4) **Non-specific voxels** (i.e., the voxels where β < 0 for print specific and speech specific contrasts). We only consider non-zero voxels. Further analyses were conducted in two steps as described below.

In the first step, we investigated, within the entire lvOT mask, whether the functional organization of lvOT voxels that responded to written and/or spoken inputs changed their category (as described above) across the two time points. To achieve this, we performed four paired t-tests to compare the number of voxels in each of the four categories between T1 and T2 corrected for multiple comparisons using Bonferroni correction. We further examined the change in lvOT’s functional organization by computing the percentages of voxels that changed their function, i.e., shifting from one category to another, from T1 to T2. The spatial distributions of the four categories of lvOT voxels at the two time points were illustrated in 3D brain maps and in a flow diagram (Figure 6). The spatial maps are based on the group averaged activation values from the 1^st^ level analysis across subjects.

In the second step, we investigated the relationship between the lvOT voxels’ functional responses and children’s reading level by computing Spearman partial correlations between the reading level measured at each time point and the number of voxels from each category, while considering age at T1and T2, sex and non-verbal IQ as covariates. Again, we conducted the same analyses excluding children with null reading scored at T1 (see SM1).

### Transparency and Openness

Behavioral and ROI data as well as the data analysis codes used in the current study are available online on the Open Science Framework data repository: https://osf.io/3t2xm/?view_only=32fe026232344e77b716d2e1cf48bae7

We reported all data exclusions, all manipulations, and all measures and statistical tools in the study in the Method section. This study’s design and its analysis were not preregistered.

For all statistical analysis and visualization, we used R software (R Core Team, 2012), IBM SPSS Statistics (Version 27) and Nilearn packages (Abraham et al., 2014).

## RESULTS

### Whole-brain analysis

Figure 1 and Table 2 showed the results from one-sample t-tests conducted on whole-brain activations at T1 and T2. The contrast between print vs. symbol strings did not show any print-specific regions at T1. At T2, the same contrast led to significant activations within the left language-related network, specifically in the lvOT, Superior Temporal Gyrus (STG), Middle Temporal Gyrus, Inferior Frontal Gyrus (IFG), Precentral Gyrus, and Supplementary Motor Area. This indicates that even in a passive viewing protocol as the one used here, young readers managed to extract linguistic information from written input. As expected, the speech vs. vocoded speech contrast revealed speech-specific regions already at T1 within the bilateral temporal network including the left Inferior Frontal Gyrus. The same pattern remained at T2. The logical conjunction between print-specific and speech-specific one-sample t-tests showed a significant print-speech convergence in the left and right posterior STG and left IFG at T2 (Fig 1., Table 2). Results from the similar analysis based on the print-rest and speech-rest contrasts are presented in SM1 (Figure S1). The main difference was the print-speech conjunction in the left IFG and bilateral STG present already at T1 due to the significant print-rest activation present at T1.

**Table 2.** One-sample t-tests at the first (T1) and second time-points (T2) for Speech-Vocoded Speech and Print-Symbols Strings contrasts as well as Conjunction analysis for the whole group (N-68) at the threshold level p<0.001, FWEc p< 0.05.

| clusters | X | Y | Z | T | Number of voxels |
| --- | --- | --- | --- | --- | --- |
| T1 |  |  |  |  |  |
| SPEECH-VOCODED SPEECH |  |  |  |  |  |
| Temporal_Mid_L (aal) | -58 | -30 | 4 | 14.8201 | 6756 |
| Temporal_Sup_R (aal) | 58 | -14 | -2 | 13.1801 | 3663 |
| Thalamus_L (aal) | -10 | -30 | 2 | 5.2266 | 146 |
| PRINT-SYMBOL STRINGS | - | - | - | - | - |
| T2 |  |  |  |  |  |
| SPEECH-VOCODED SPEECH |  |  |  |  |  |
| Temporal_Inf_R (aal) | 40 | 0 | -40 | 5.1078 | 101 |
| Temporal_Inf_L (aal) | -38 | 0 | -40 | 5.8062 | 198 |
| Temporal_Sup_L (aal) | -60 | -8 | -2 | 18.1837 | 4860 |
| Temporal_Mid_R (aal) | 62 | -8 | -4 | 16.1014 | 3944 |
| Thalamus_L (aal) | -8 | -28 | 4 | 5.1118 | 69 |
| Thalamus_R (aal) | 12 | -26 | 0 | 4.5046 | 54 |
| Frontal_Inf_Oper_L (aal) | -46 | 16 | 22 | 4.9006 | 125 |
| PRINT-SYMBOL STRINGS |  |  |  |  |  |
| Temporal_Inf_L (aal), Fusiform_L (aal) | -42 | -48 | -14 | 5.6151 | 99 |
| Insula_L (aal), Frontal_Inf_Tri_L (aal),<br>Frontal_Inf_Orb_L (aal),<br>Temporal_Pole_Sup_L (aal) | -32 | 24 | 4 | 5.8868 | 233 |
| Temporal_Sup_L (aal), Temporal_Mid_L (aal) | -58 | -2 | -8 | 4.0696 | 23 |
| Temporal_Mid_L (aal), Temporal_Sup_L (aal),<br>SupraMarginal_L (aal) | -54 | -42 | 6 | 7.3301 | 969 |
| Frontal_Inf_Oper_L (aal), Frontal_Inf_Tri_L<br>(aal), Precentral_L (aal), Rolandic_Oper_L (aal) | -48 | 16 | 24 | 5.4349 | 382 |
| Temporal_Mid_R (aal), Temporal_Sup_R (aal),<br>SupraMarginal_R (aal) | 50 | -38 | 6 | 5.0857 | 251 |
| Insula_R (aal), Frontal_Inf_Tri_R (aal) | 36 | 26 | 2 | 5.3231 | 71 |
| Postcentral_L (aal) | -60 | -18 | 26 | 3.9025 | 20 |
| Precentral_L (aal), Postcentral_L (aal) | -52 | -2 | 48 | 7.2604 | 221 |
| Supp_Motor_Area_L (aal),<br>Supp_Motor_Area_R (aal), Cingulum_Mid_L<br>(aal), Cingulum_Mid_R (aal),<br>Frontal_Sup_Medial_L (aal),<br>Frontal_Sup_Medial_R (aal) | -4 | 8 | 56 | 7.5862 | 557 |
| CONJUNCTION PRINT & SPEECH |  |  |  |  |  |
| Temporal_Sup_L (aal) | -62 | -28 | 4 | 13.602 | 920 |
| Temporal_Sup_R (aal) | 50 | -32 | 4 | 10.9888 | 234 |
| Frontal_Inf_Oper_L (aal) | -46 | 16 | 22 | 4.9006 | 79 |

### The lvOT analysis

Figure 3 shows a developmental increase in the print-specific as well as speech-specific activation within the lvOT mask. The 2×2 repeated measure ANOVA showed a significant main effect of type of contrast (F=697, p<0.001), time point (F=546, p<0.001), and of type of contrast * time point interaction (F=5.5, p<0.01). Further analyses of the time point effect for each type of contrast showed that, two years after the onset of formal reading instruction, the print-specific activation significantly increased (p<0.001) and the speech-specific activation shifted from a negative value to a positive value (p<0.001).

**Figure 3.**
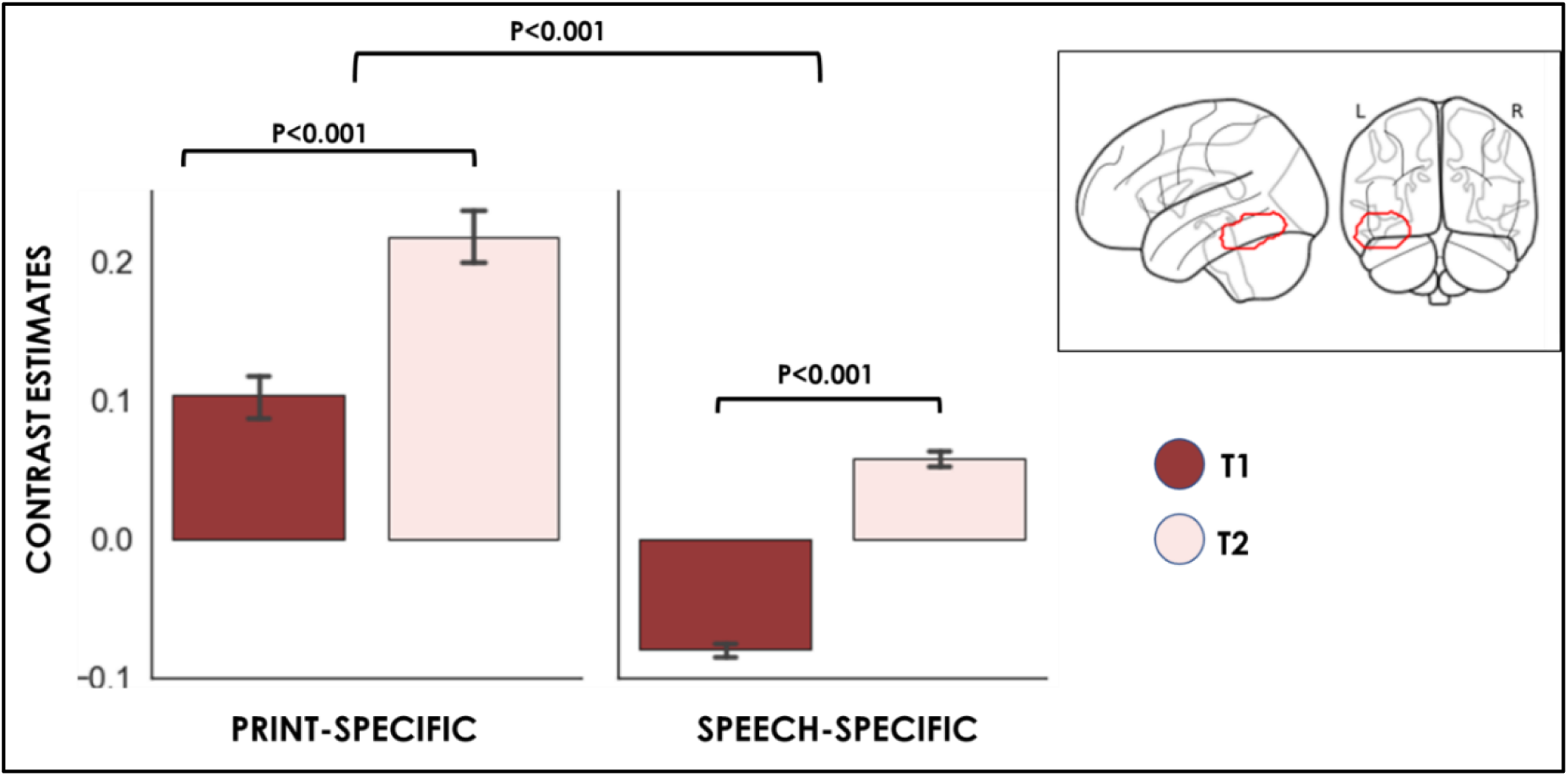
Pattern of print-specific and speech-specific activation within the lvOT mask (2189 voxels) observed across the two time points.

The investigation of the relationship between reading level (number of words read per minute) and the activation elicited by the print-specific and speech-specific contrasts within lvOT mask with Age, Sex, Non-verbal IQ as covariates showed that there was a significant increase of lvOT’s print-specific activity with reading score at T1 (265 voxels, peak coordinates: −40 −52 - 16, T=5.32). No such relationship was observed at T2. No significant correlation was observed between reading expertise and speech-specific activity at either time point. The same result pattern was also observed in the additional ROI analyses where the data from the 12 children with null reading score at T1 were excluded (236 voxels, T=4.87, SM1).

As shown in Figure 4, the distinct patterns observed at T1 and T2 could be explained by a gradual increase of lvOT activation specific to printed words at the initial stage of reading acquisition (red dots). From a certain level of reading expertise, the level of lvOT activation became relatively stable, at least in the task used here (pink dots).

**Figure 4.**
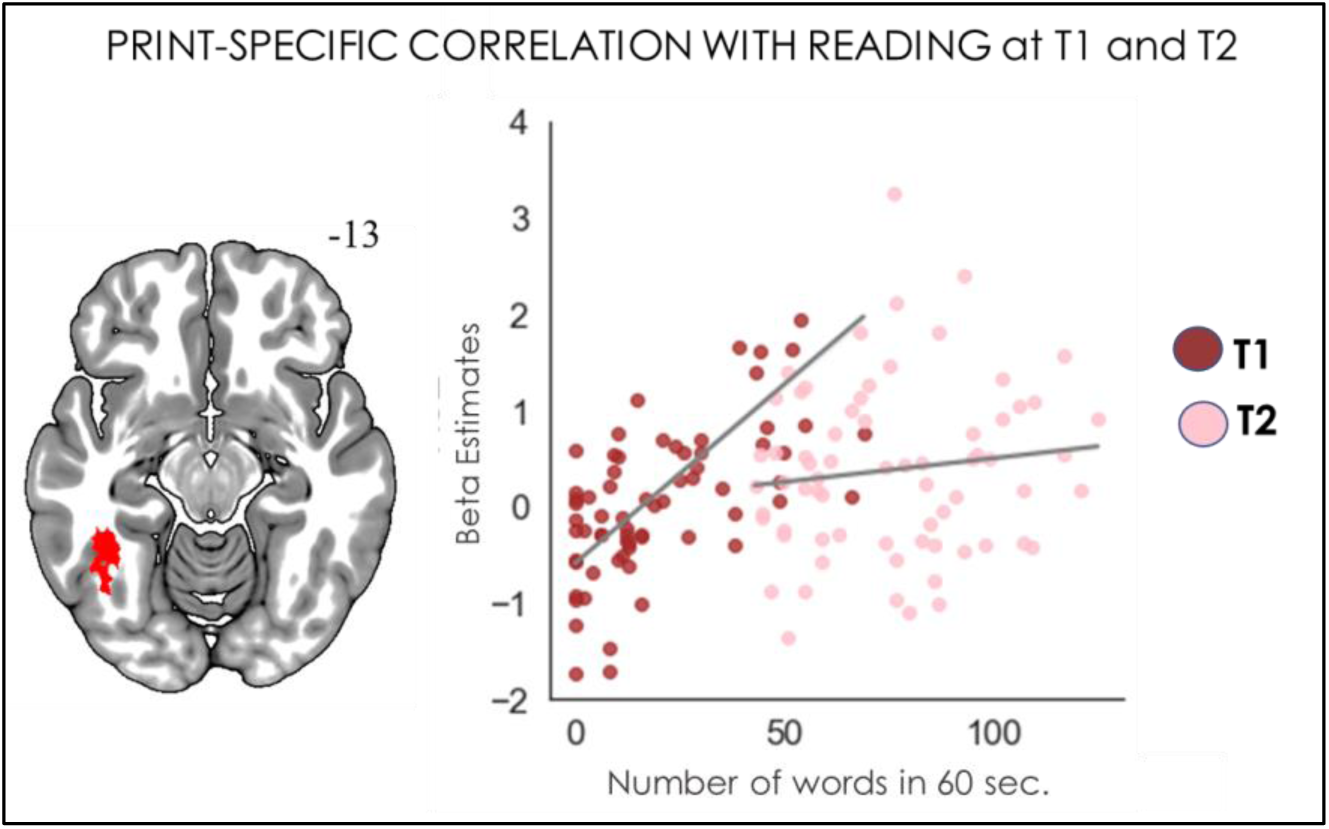
Cluster showing a significant correlation between reading score at T1 and the level of activation for the print-specific contrast within lvOt at T1 and T2.

### Individual voxels analysis within the lvOT

Based on an anatomical lvOT mask that contained 2189 voxels, we computed the number of voxels that showed the print-speech specific, print-specific, speech-specific, and non-specific response patterns.

Paired t-tests with the Bonferroni correction (adjusted threshold: p<0.0125) showed that the number of print-speech specific convergent voxels significantly increased from T1 to T2 (T1 = 492 vs. T2 = 633, p<0.012). The number of Print-Specific, Non-specific and Speech-Specific voxels did not change significantly between the two time points (Print-Specific = 595 (T1) vs 495 (T2), p=0.09, Non-specific = 588 vs 538 p=0.41, Speech-Specific = 507 vs. 516 voxels, p=0.88).

At T1, the lvOT mask showed that 22.5% of the voxels were print-speech specific, 27.2% were print-specific, 26.9% were non-specific, and 23.2% were speech-specific. At T2, the percentage of print-speech specific voxels increased to 28.9%, while print-specific voxels decreased to 22.6%, non-specific voxels to 24.6%, and speech-specific voxels remained relatively stable at 23.6%.

**Figure 5.**
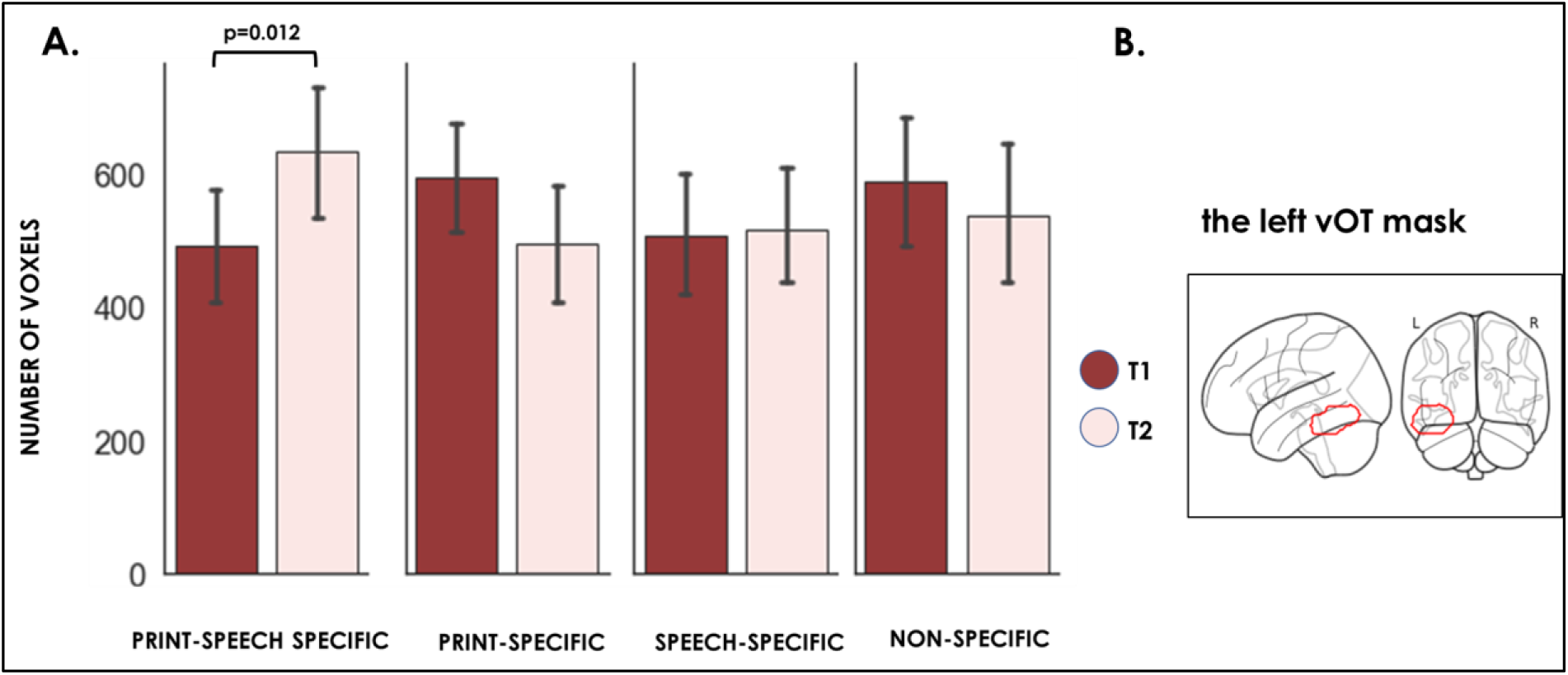
The number of voxels in the print-speech specific, print-specific, speech-specific and non-specific category within the lvOT mask at the two time points. Plots based on the activity from the individual first-level SPM maps.

A more detailed illustration of the changes in lvOT functional organization is illustrated in the distribution flow in Figure 6, based on the group averaged data calculation. It showed that for all the print-speech specific voxels observed at T2, 54% were print-specific voxels, 29% were print-speech specific voxels, 3% were speech-specific voxels, and 12% were non-specific voxels at T1. Figure 6C illustrates the spatial localization of the different categories of voxels within the lvOT. Interestingly, the print-specific voxels at T1 that gained their additional specificity to speech sounds at T2 were mainly located at the most anterior medial part of the lvOT.

**Figure 6.**
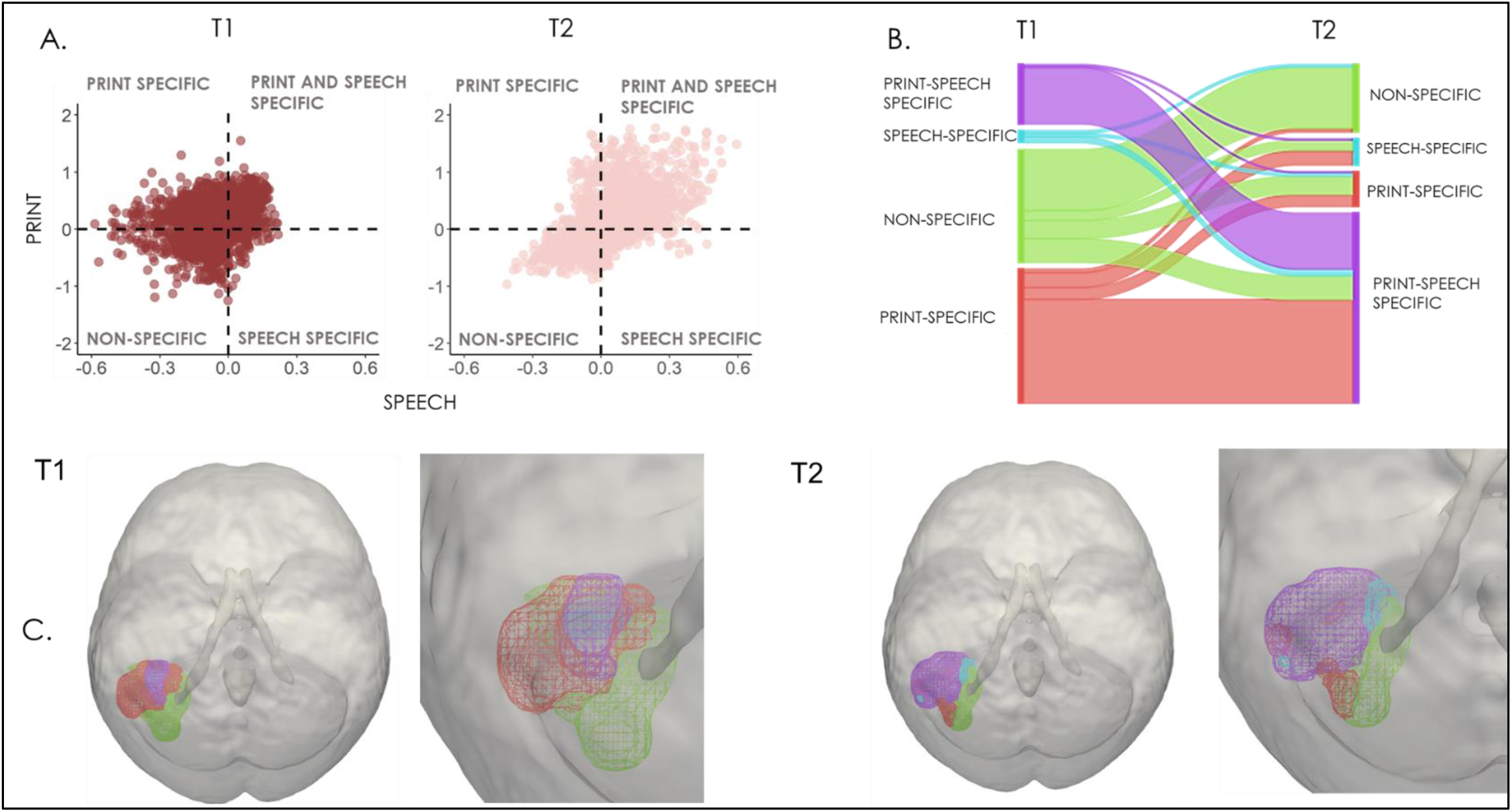
A.) The distribution of 2189 lvOT voxels activation in the four categories: print-speech convergent, print-specific, speech-specific activations and non-specific across the two time points. B) A flow chart illustrating the changes in voxel category across the two time points. C) Localization maps of the voxel category distribution. Plots based on the group averaged data values.

In the final analyses, we investigated whether the distribution of the functionally defined voxels was related to the reading skill. Spearman partial correlations between the number of voxels from each category and reading score (number of words read correctly per 60 sec.) at each time point were conducted. Age, sex and non-verbal IQ were considered as covariates. The analyses showed that the number of print-speech specific voxels was positively related to the reading level at T1 (Rho ρ = 0.4, p< 0.001) while the same correlation became non-significant at T2. Also, we observed a negative correlation between the number of Non-specific voxels and the reading level at T1 (Rho ρ = −0.3, P<0.005, See Figure 7). The results were replicated after excluding the 12 children who showed a null score in the reading task (Print-Speech-reading, Rho ρ=0.2, p<0.05, Non-specific, Rho ρ = −0.2, p<0.05, See Figure S6). No other comparisons survived the Holm-Bonferroni correction for multiple testing.

**Figure 7.**
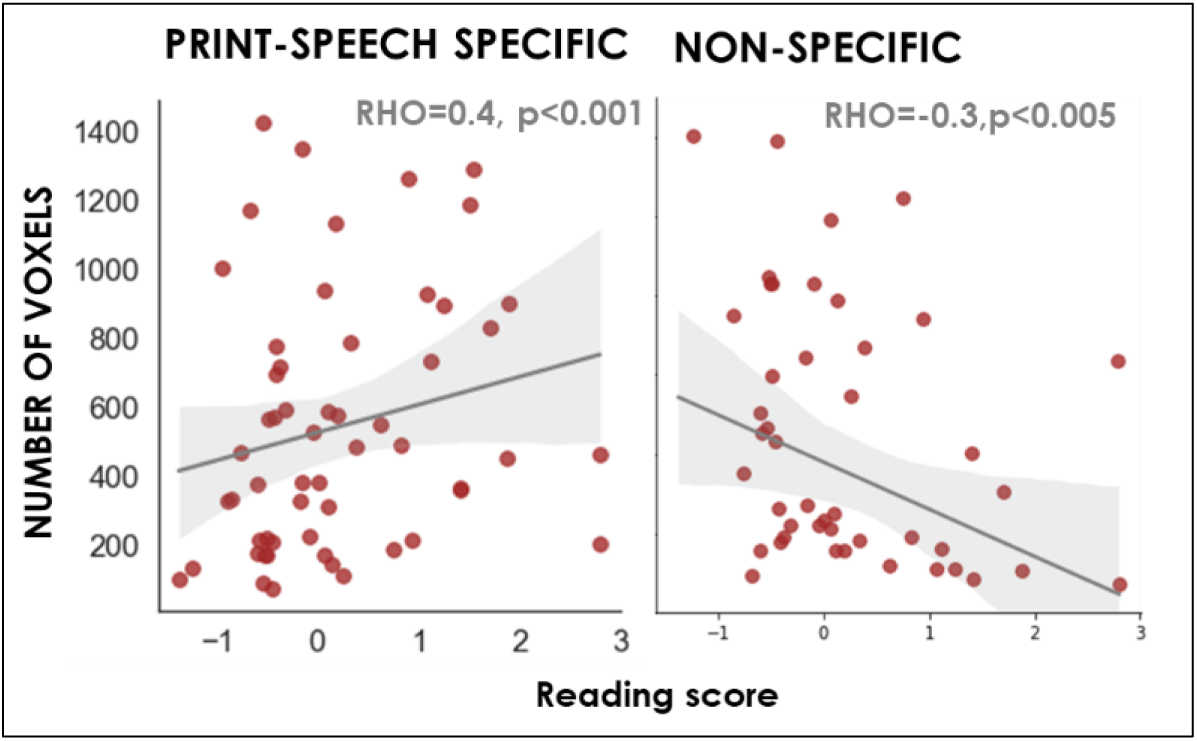
Significant partial correlations between the number of voxels in print-speech specific and non-specific categories and the reading score T1, after controlling for the Age, Sex and Non-verbal IQ.

**Table 3.**
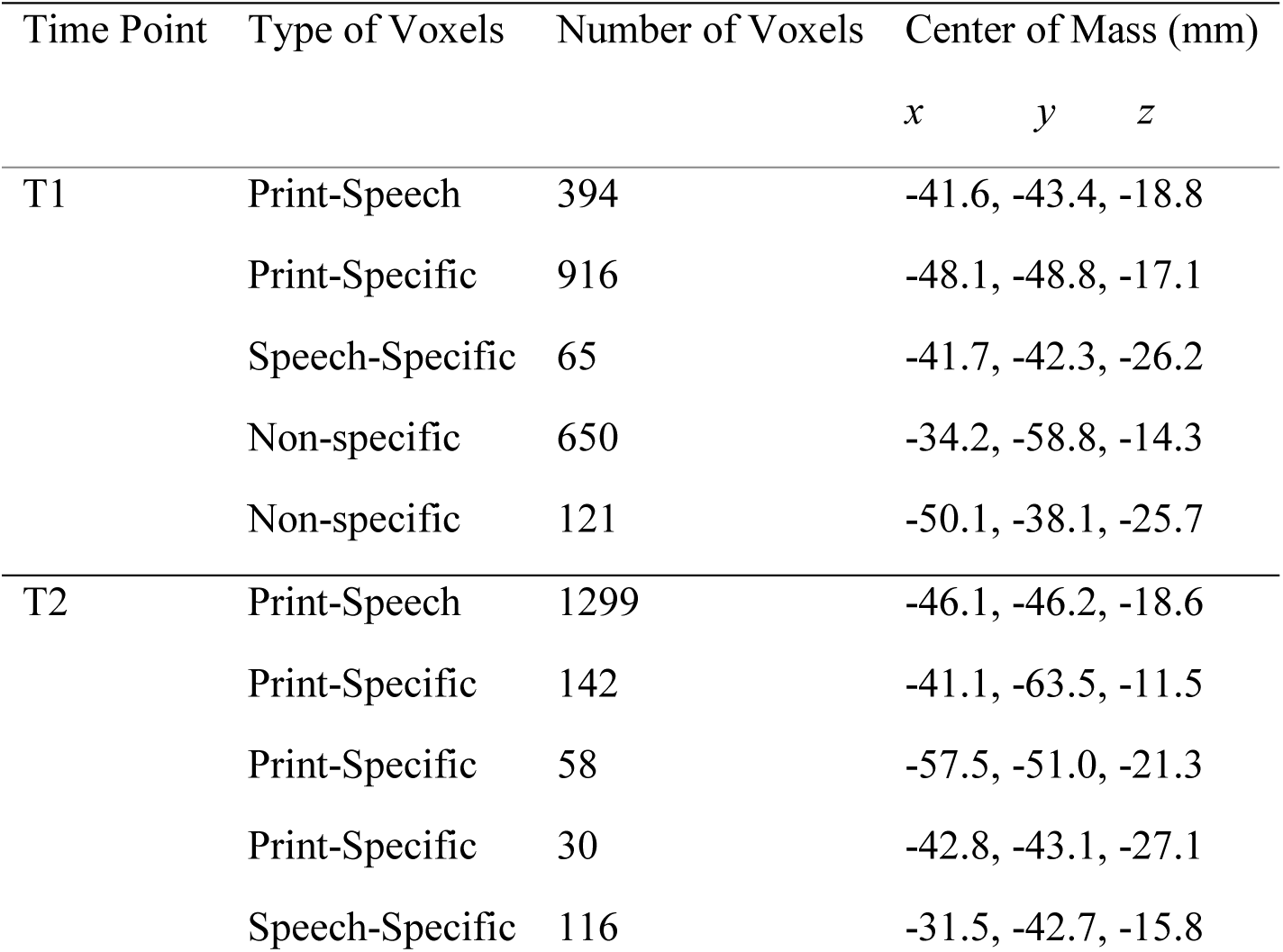
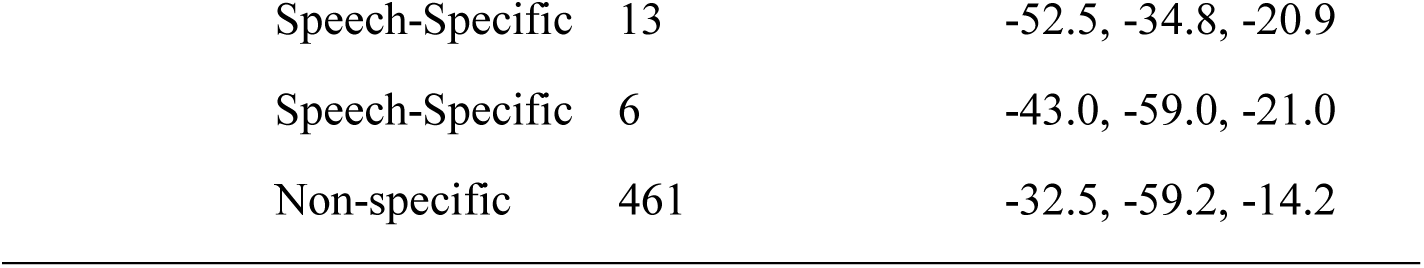
The localization of the voxels in lvOT from the four voxel categories at the two time points.

| Time Point | Type of Voxels | Number of Voxels | Center of Mass (mm) |  |  |
| --- | --- | --- | --- | --- | --- |
|  |  |  | *x* | *y* | *z* |
| T1 | Print-Speech | 394 | -41.6, | -43.4, | -18.8 |
|  | Print-Specific | 916 | -48.1, | -48.8, | -17.1 |
|  | Speech-Specific | 65 | -41.7, | -42.3, | -26.2 |
|  | Non-specific | 650 | -34.2, | -58.8, | -14.3 |
|  | Non-specific | 121 | -50.1, | -38.1, | -25.7 |
| T2 | Print-Speech | 1299 | -46.1, | -46.2, | -18.6 |
|  | Print-Specific | 142 | -41.1, | -63.5, | -11.5 |
|  | Print-Specific | 58 | -57.5, | -51.0, | -21.3 |
|  | Print-Specific | 30 | -42.8, | -43.1, | -27.1 |
|  | Speech-Specific | 116 | -31.5, | -42.7, | -15.8 |

|  |  |  |
| --- | --- | --- |
| Speech-Specific | 13 | -52.5, -34.8, -20.9 |
| Speech-Specific | 6 | -43.0, -59.0, -21.0 |
| Non-specific | 461 | -32.5, -59.2, -14.2 |

## DISCUSSION

Our longitudinal study tracked young readers from the beginning of formal reading instruction to the second or third grade of elementary school. The results obtained in the analyses conducted at three spatial scales suggest a functional reorganization of neural responses within the left ventral occipitotemporal cortex, which reflects print-speech specific convergence and its relationship with the reading level.

### Whole-brain level

The results from the whole-brain analysis revealed that brain areas showing speech-specific activation were already present at T1 and remained stable until T2. In contrast, areas showing print-specific activation only emerged at T2. The areas included not only the lvOT but also several areas that are involved in the phonological, semantic and articulatory components of the spoken language network, i.e., STG, MTG, IFG, Precentral Gyrus, and Supplementary Motor Area. Such high-level processing of written input was not observed at T1, which could be explained by the fact that, at an early stage of reading acquisition, the lvOT’s sensitivity to print versus symbol strings is still developing and might have strong variability across subjects. This interpretation is with line with the presence lvOT activation already at T1 when activation to print input was contrasted against baseline.

Regarding our core interest in the emergence of print-speech specific convergence response, the conjunction analysis only showed a significant activation at T2 in bilateral pSTG and left IFG regions. Importantly, no convergence response was found in the lvOT at any time point. This observation replicates the previously reported early print-speech specific convergence response occurring within the spoken language network to accommodate print acquisition and a delayed, or absence of, such response in the visual ventral pathway in transparent languages, like Polish (Rueckl, 2015; Chyl et al., 2018, 2021). However, as discussed below, more detailed analyses at the voxel level allowed us to deepen our investigation on the functional reorganization within the lvOT and revise the conclusion obtained at the whole brain level.

### The left-vOT level

The analyses focusing on the lvOT provided evidence that the print-specific and speech-specific responses significantly increased from T1 to T2. Interestingly, the speech-specific activation, which was below the baseline level, thus indicating a cross-modal suppression pattern of speech-specific processing (Baier et al., 2006) at the initial stage of reading acquisition (T1), became positive at T2. This modulation of cross-modal responses suggests that increasing reading experience renders the lvOT sensitive to spoken language.

The response to print-specific stimuli within lvOT was correlated with the reading level at T1 although the correlation did not reach significance at T2. An improvement in reading ability could explain this change across the two time points during the two years. As predicted by *Interactive Activation account* and the *Predictive Coding framework* proposed by Price and Devlin (2011, see also Brem et al. 2010, Yeatman, et al., 2012), the relationship between the strength of lvOT response to print and reading ability would have an inverted-U shape across different stages of reading acquisition. At the prereader stage, the lvOT activity would be driven only by visual input, given the absence top-down knowledge associated with the input. This pattern would be followed by a boost of lvOT activity at the initial stage of reading acquisition when children learn to associate visual symbols with sounds and meanings, although the integration between bottom-up and top-down information is still effortful and error-prone. At the final stage, when reading becomes more automatic, brain activation is initially sustained, then decreases compared to earlier stages, due to more efficient integration of bottom-up and top-down information and reduced prediction errors. In the present study, at T1, children were at the initial stage of reading acquisition and we indeed observed a linear relationship between brain activity and reading expertise. At T2, after two years of reading instruction, given the nature of low-level passive viewing task that we used to reveal lvOT activation, the highest and stable activation level was reached, which could explain the absence of its correlation with reading score. In line with our observation, a recent longitudinal fMRI study conducted on 16 children who were typical and poor readers showed a gradual increase in lvOT activation to letters at the start of primary school, followed by a plateau around the 2^nd^ and 3^rd^ Grade and then a decline of activity around the 4^th^ Grade (Di Pietro et al., 2023).

Finally, we did not find a correlation between speech-specific activity in lvOT and reading level at T1 nor T2. Again, this could be due to the nature of the passive task used to show speech-specific activation. Former studies already showed that lvOT response is typically low or non-significant in such tasks even in skilled readers (Dehaene & Cohen, 2010; Planton et al., 2019). While a low-level passive viewing task seems suitable to reveal brain responses in young children who have just begun reading acquisition, it is interesting to investigate in future studies, whether a different developmental trend emerges in more demanding tasks, especially in participants with higher reading skills. An influence of task demands on longitudinal changes in lvOT was indeed recently reported by Ozernov-Palchik et al. (2023).

### Individual lvOT voxels level

So far, the classic analyses at the whole brain level failed to reveal print-speech specific convergence response in the lvOT, which replicated previous findings in transparent writing systems (Rueckl et al., 2015, Chyl et al., 2018, 2021, Dębska et al., 2021). In the final set of analyses, we proposed an original way to explore the data by looking at the developmental trajectory of the functional reorganization of the lvOT responses to print-speech convergence at the voxel level. To this aim, we separated the lvOT voxels into four categories according to their response profiles to print and speech specific input, i.e., those that showed print-specific responses, those that showed speech-specific responses, those that showed both print- and speech-specific responses, and those that showed no-specific response to either language input.

Based on the comparisons of the number of voxels from the different categories across the two time points we found a significant increase in the number of lvOT voxels that jointly responded to both print and speech, and stable number of voxels that showed speech-specific, non-specific and print-specific responses, although a slight decline was observed in the last category. A further examination of these developmental changes in voxel category showed that the increase in the number of voxels that responded to both print and speech specific stimuli, thus reflecting its convergence, was mainly due to the transformation of print-specific to print- and speech-specific voxels. Indeed, the majority (54%) of print- and speech-specific voxels at T2 were print-specific at T1. To our knowledge, this is the first time that a voxel-based analysis was used to reveal a shift in functional responses in lvOT in the context of the reading acquisition.

We also found that at T1, reading score was positively correlated with the number of print-speech specific voxels and negatively correlated with the number of non-specific voxels. This is in line with our hypothesis that the increase of cross-modal sensitivity would be positively related to the reading level and not necessarily the mere increase in specific responses to a single modality. No other significant correlations were reported. In line with the results obtained at the ROI level, the significant correlations were restricted to the initial stage of reading acquisition, which could partly be to the low-level passive viewing and listening tasks used in the scanner.

## Conclusions

Previous studies showed that print-speech convergence within the higher-level visual cortex was less consistent that the one observed in spoken language regions like the pSTG (Chyl et al., 2021, Debska et al., 2021). Our initial analysis at the whole-brain level replicates these observations. However, a more fine-grained examination of brain responses at the voxel level revealed novel insightful information which suggests that, even in a transparent writing system, learning to read and improving reading skills not only lead to an overall increase of lvOT sensitivity to each of the two language modalities but also to their convergence. We found that the initial print specialization regions developed sensitivity to speech sounds with reading experience. The convergence responses were positively related to the reading skill at the earliest stage of literacy acquisition. Such cross-modal functional reorganization occurs in both the ventral visual pathway and in the spoken language system as reported in the literature and our study (see: Methods & Results, SM3). Overall, this increase of convergence between different modalities of language representations through reading acquisition illustrates the plasticity of the human brain to accommodate the acquisition of new skills.

## ACKNOWLEDGMENTS

This work was supported by the French Ministry of Research: ANR-19-CE28-0001-01 (to C.P.), ANR-16-CONV-0002 (ILCB), ANR-11-LABX-0036 (BLRI) and the Excellence Initiative of Aix-Marseille University (A*MIDEX). Centre de Calcul Intensif d’Aix-Marseille is acknowledged for granting access to its high-performance computing resources.

## List of Supplementary Materials

S1. One-sample t-tests for print-rest and speech-rest activation at the first and second time-point and logical conjunction results.

S2. Behavioral and brain correlation with the reading level on the readers group (N=56) after excluding the prereaders group, i.e., children who obtained 0 in the word reading task at T1 (N=12).

S3. Individual voxels analysis within the left posterior superior temporal gyrus: Method & Results.

## Supplementary Materials S1

**Figure S1.**
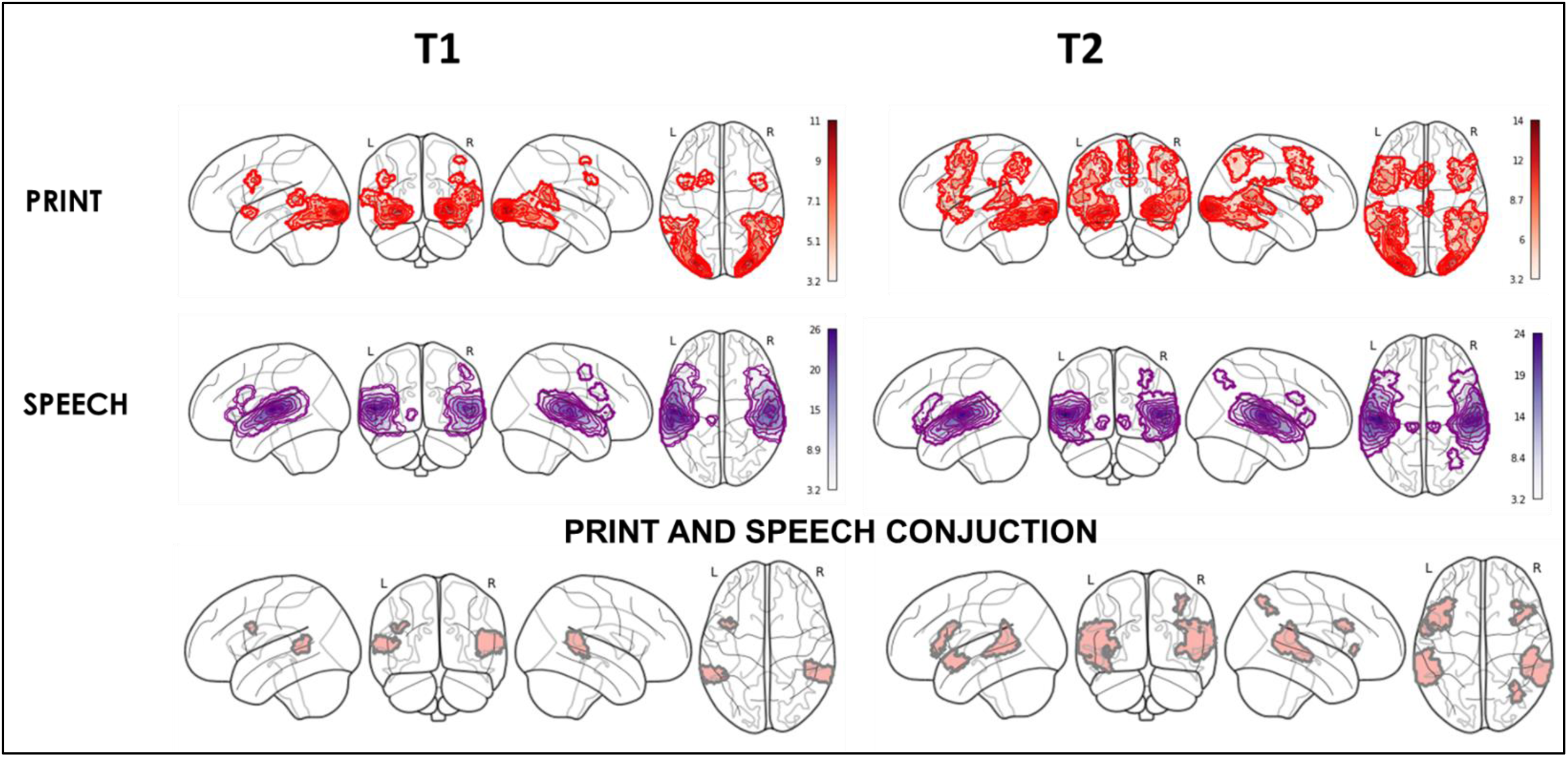
One-sample t-tests for print-rest (red) and speech-rest (purple) activation at the first (T1) and second time-point (T2) and logical conjunction results at T1 and T2.

## Supplementary Materials S2

Behavioral and brain correlations with the reading level in the readers group (N=56), after excluding children who obtained 0 in the word reading task at T1 (N=12)

**Table S1.** Averaged performances obtained in the behavioral tasks by the 12 prereaders. Standard Deviations are in the parentheses.

|  | Non-verbal_IQ | Age_tp1 | Age_tp2 | Letter_tp1** | Word_tp1*** | Orthography task TP1# | Word_tp2 |
| --- | --- | --- | --- | --- | --- | --- | --- |
| Mean | 78 (25) | 6.56(0.4) | 8.6 (0.4) | 25 (11) | 0 | 19(3.9) | 66 (15) |
| Min | 24 | 5.97 | 7.8 | 4 | 0 | 16 | 45 |
| Max | 97 | 7.6 | 9.1 | 47 | 0 | 26 | 93 |

\*Standardized score, Raven Matrices Test, measured only at T1; \*\* Knowledge of letters, max. 64 (upper and lower cases); \*\*\* Number of words read correctly in 60 seconds # max. 30.

**Figure S2.**
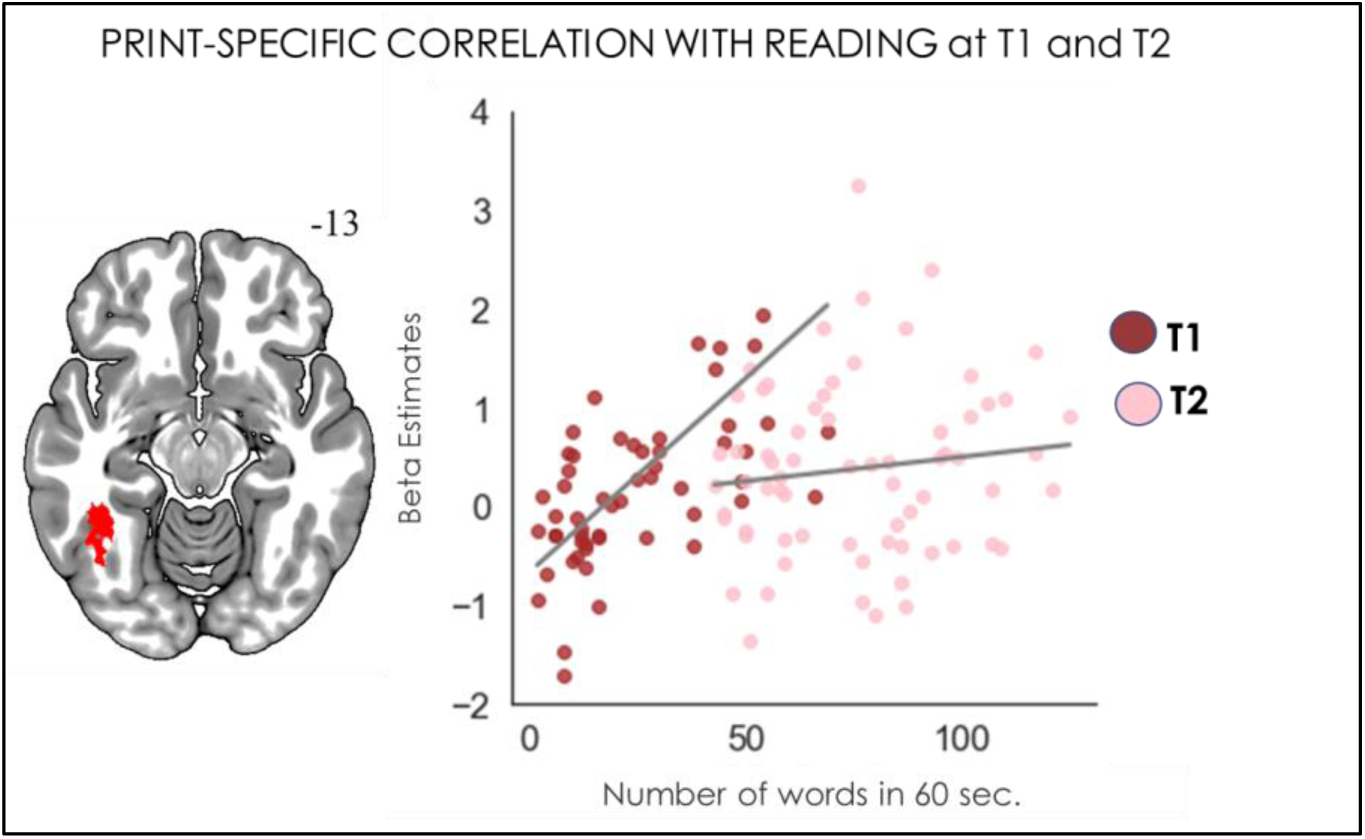
Regression model of lvOt cluster with the reading level (N=56) after excluding prereaders. Cluster showing a significant correlation between reading score at T1 and the level of activation for the print-specific contrast within lvOt at T1 and T2.

**Figure S3.**
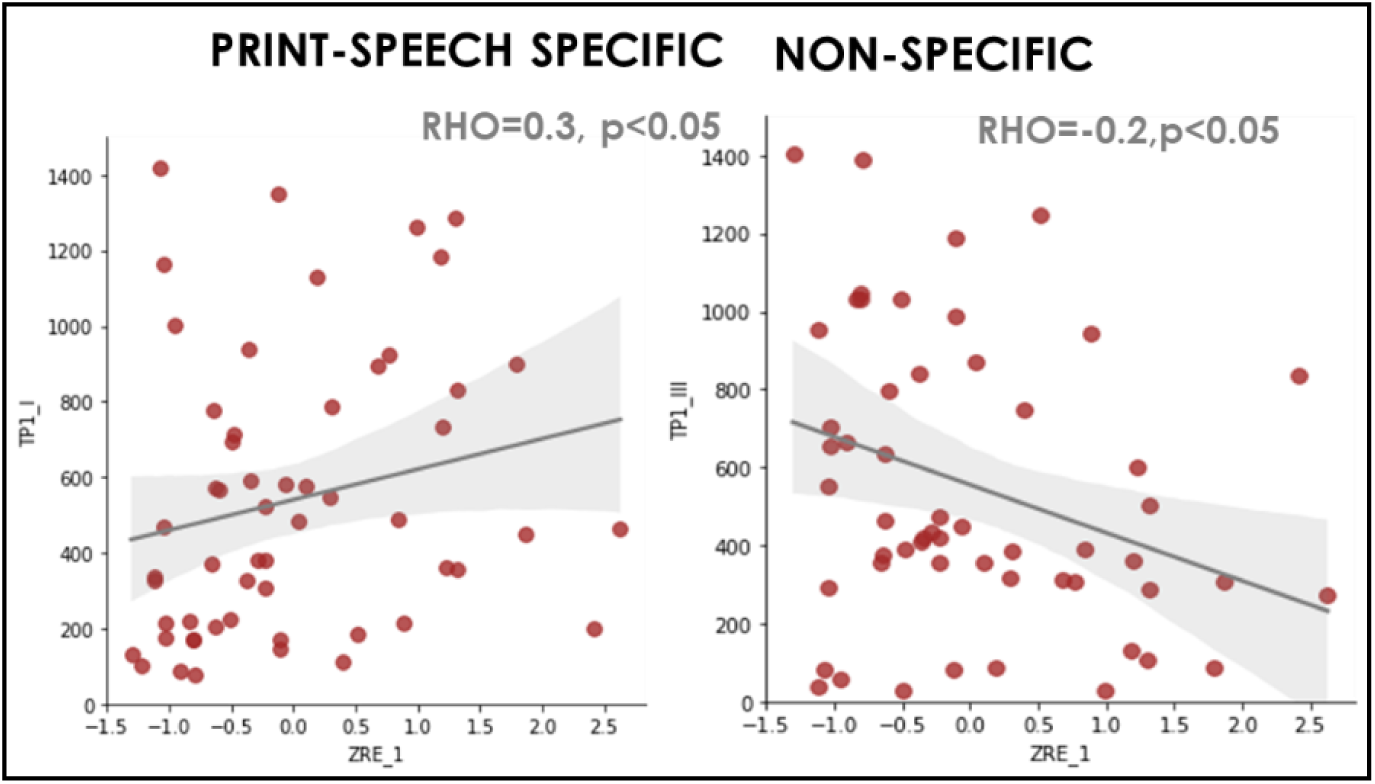
Significant correlations between the number of voxels in print-speech specific and non-specific categories and the reading score T1, after controlling for the Age, Sex and Non-verbal IQ exluding 12 prereaders.

## Supplementary Material S3

The left posterior superior temporal gyrus voxel analysis: methods & results.

### Methods

**Figure S4.**
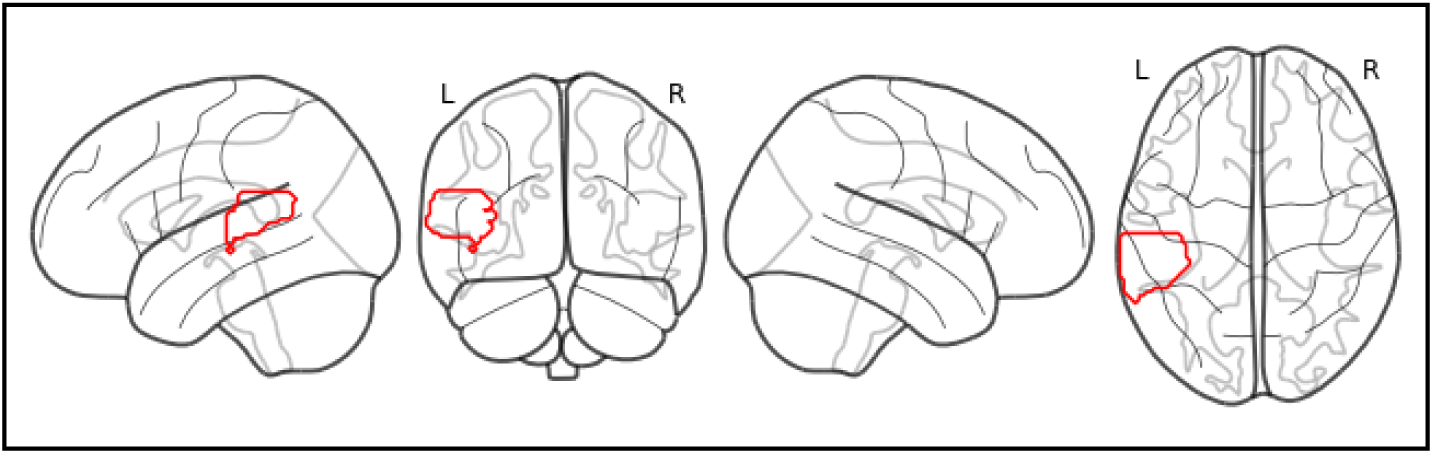
The lpSTG mask contains 1210 voxels encompassing the posterior part of the Superior Temporal Gyrus, based on an anatomical template from AAL atlas).

Within the left pSTG mask (1210 voxels), based on the 1^st^ level activation values for each participant, we defined four categories of voxels in T1 and T2: 1) **Print- and Speech-specific voxels** (i.e., the voxels active above threshold where beta value β > 0 in both print-specific and speech-specific contrasts; 2) **Print-specific voxels** (i.e., the voxels active above threshold beta value β > 0 in print specific contrast and β < 0 in speech specific contrast); 3) **Speech-specific voxels** (i.e., the voxels active above threshold beta value β > 0 in speech specific contrast and β < 0 in print specific contrast); 4) **Non-specific voxels** (i.e., the voxels where β < 0 for print specific and speech specific contrasts). We only consider non-zero voxels. Localization plots S5. were computed based on group-averaged data values.

### Results

**Figure S5.**
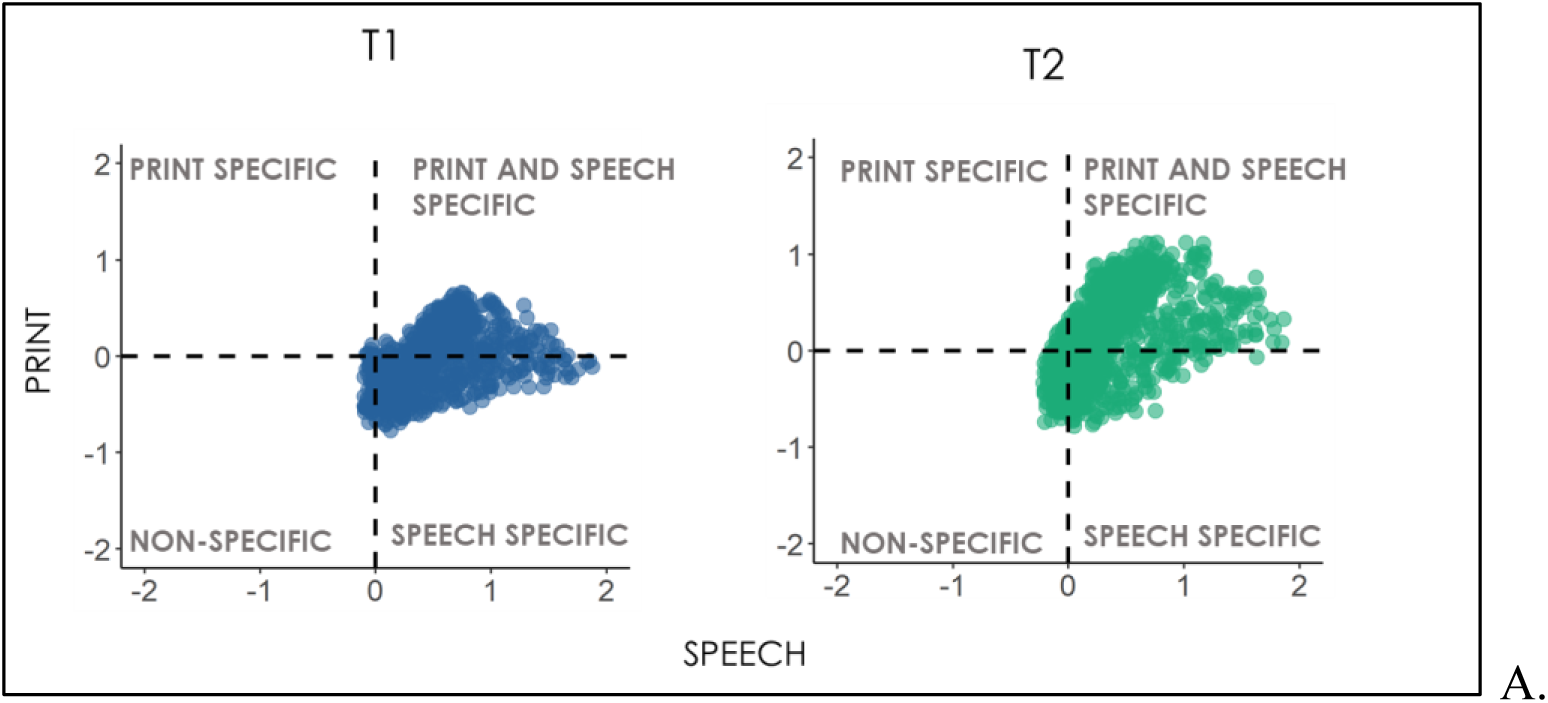
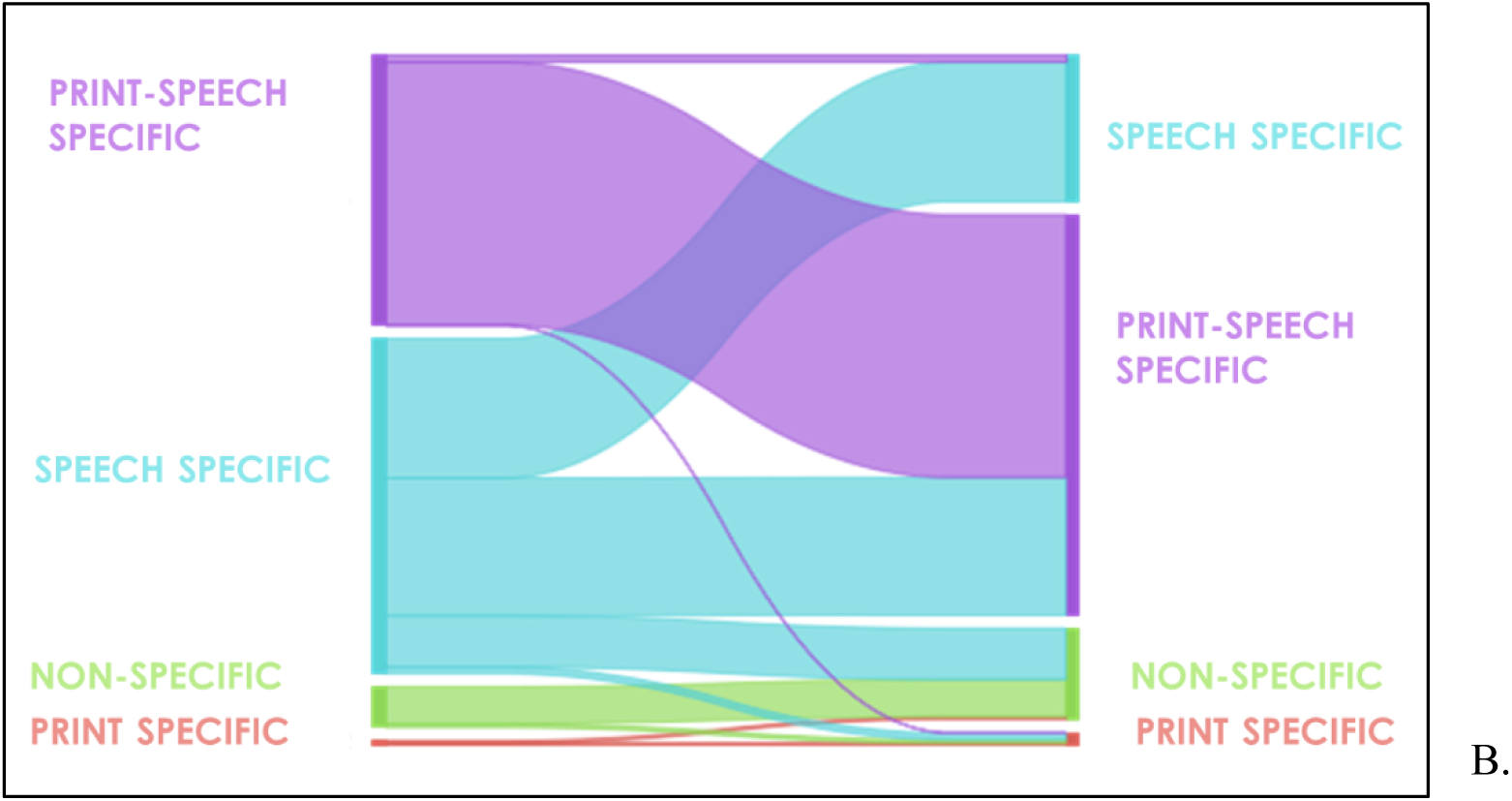
The distribution (A) and flow plot (B) of voxel categories between T1 and T2.

**Figure S6.**
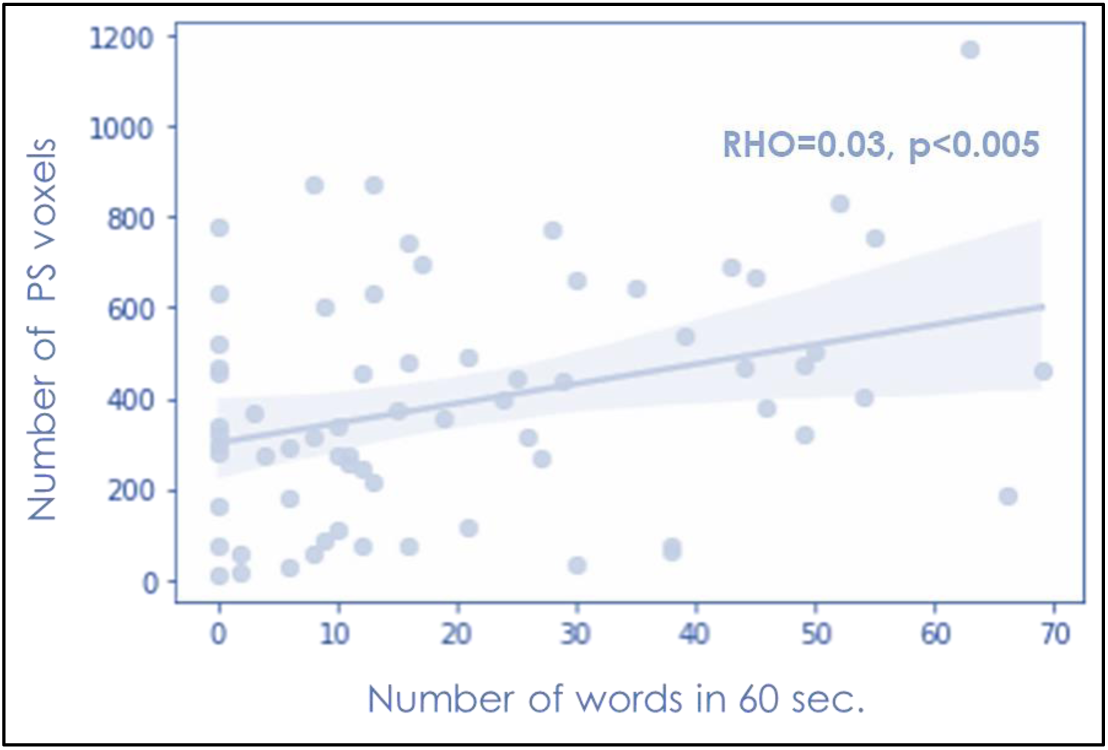
The significant correlation of the number of print-speech specific voxels in pSTG and the reading level at T1. Rho = 0.32, p<0.005.

